# Using Community Science and eDNA to Inform a State-Designed Marine Protected Area Monitoring Program: A Case Study from Malibu, California

**DOI:** 10.64898/2026.09.29.755491

**Authors:** Tanner Waters, Zachary Gold, Emily Parker, Olivia Simon, Liam Jenny, Richard F. Ambrose, Paul H. Barber, Robert A. Eagle

**Author notes:** Address for correspondence: Tanner Waters; Robert Eagle.

## Abstract

Management of marine protected areas (MPAs) require long-term monitoring and evaluation that is often limited by the cost and effort of conventional visual based surveys. We evaluated a community science-based eDNA approach to MPA monitoring over two years in the Point Dume State Marine Reserve, the Point Dume State Marine Conservation Area, and two paired unprotected reference sites. Across the sampling period, we detected 119 species of fish and marine mammals, including 19 of the 22 regional management-priority species for Southern CA MPAs. Protected and unprotected comparison sites were broadly similar in overall assemblage with 93 species shared across all sites. Multivariate analyses revealed site-level differences, although season and year explained more variation than protection status. Species presence also shifted during a La-Niña transition, highlighting the importance of long-term monitoring under different climatological contexts. Community scientists greatly expanded the temporal and spatial scope of eDNA sampling and generated high-quality data. Subsampling analysis showed that quarterly sampling, with three sample replicates and sequencing depths between 700k-990k, recovered 95% of observed biodiversity of the full time series. These results demonstrate that a community science-based eDNA program could provide a standardized, scalable, low-cost, and management-relevant approach for long-term monitoring across California’s MPA network.

## Introduction

Marine protected areas (MPAs) are critical tools for preserving biodiversity and mitigating the impacts of overfishing and other human pressures on coastal ecosystems^1^. By restricting extractive activities, MPAs are designed to protect species diversity and rebuild depleted fish populations^2,3^. However, demonstrating that MPAs are achieving their intended goals is important for justifying the added limitations that these areas impose on human-use activities as well as guiding adaptive management strategies and informing future MPA development^4^. This requires regular, long-term biological monitoring that can reliably detect changes in community composition and species occurrence through time^5^.

The need for effective MPA monitoring is particularly important in the Southern California Bight, a region which contains highly productive reef and kelp forest ecosystems. This region supports more than 500 species of fish^6^, underscoring the ecological, economic, and social value of Southern California’s ocean ecosystems. These systems are increasingly being threatened by human activities, particularly overfishing, which poses one of the largest threats to fish populations^7,8^. In response to these pressures, California enacted the Marine Life Protection Act in 1999, establishing the first statewide network of MPAs in the United States.

The California MPA network was fully implemented in 2012 and comprises 124 MPAs protecting approximately 15% of the State’s waters, with 50 MPAs in the Southern California Region alone. These MPAs vary in design and regulations, ranging from fully protected no-take State Marine Reserves to more permissive State Marine Conservation Areas that allow for some commercial and recreational take^9^. Baseline and continued monitoring of California’s MPAs have documented positive biological responses in some regions, particularly with increases in fish biomass, density, and species richness within protected areas^10–13^. However, current State MPA monitoring efforts only cover a small fraction of MPAs and habitat types within the MPA network^14,15^ because of the physical and economic challenges of sustained long-term monitoring across a spatially expansive MPA network. These challenges are likely to intensify as California advances its “30x30” initiative, which aims to conserve 30% of state lands and waters following a 2020 executive order signed by Governor Gavin Newsom and subsequent legislation in 2021. Thus, expanding the spatial extent of MPA’s will increase the need for scalable monitoring of these MPAs and their effectiveness.

A major constraint on long-term and large-scale MPA monitoring is the reliance on traditional scuba-based visual surveys for species identification. Although a well-established method for rocky reef biodiversity monitoring, scuba surveys are logistically complex in the field, resource intensive, and field-conditions dependent-limiting the scope and scale of MPA monitoring efforts^16^. Additionally, scuba surveys largely focus on a limited number of indicator species (i.e., species that are either commercially valuable, easy to measure, sensitive to human impacts, or are strong ecological drivers) rather than whole ecosystem-wide surveys^17^. Recently, California allocated $9.5 million for seven projects to expand long-term MPA monitoring^18^. However, this funding explicitly prioritizes Tier I MPA sites, with Tier II and Tier III sites monitored as capacity and budget allow. Given that there are only 34 Tier I MPAs, 72.6% of the State’s MPAs remain unmonitored, limiting the ability to detect community changes or establish site-specific baselines.

A newer, alternative approach to scuba-based monitoring is the use of environmental DNA (eDNA) metabarcoding. By detecting DNA shed by organisms in the ocean, eDNA facilitates broad taxonomic surveys from a relatively small number of seawater samples. Numerous studies show that eDNA matches or outperforms traditional visual surveys in identifying species presence or absence, including rare or cryptic taxa^19^. Moreover, the simplicity of sweater sampling relative to scuba surveys makes eDNA metabarcoding a much more cost effective and conditions-independent tool for monitoring^20^. These characteristics make eDNA particularly attractive for regular, long-term MPA monitoring.

Importantly, the logistical simplicity of eDNA sampling (i.e., sampling a liter of water and allowing it to drip through a filter) enables easy integration with community scientists, which has become an increasingly important part of coastal monitoring within California^21^. Programs like MPA Watch engage volunteers to document human use patterns and compliance within MPAs^22^, but are not currently designed to survey biodiversity. Community science-led eDNA sampling allows for volunteers to collect samples that can be developed into high-quality biological data with minimal field equipment and training^23,24^. Forgoing the need for highly skilled field-based scientists, eDNA biodiversity surveys can greatly expand the spatial and temporal resolution of MPA monitoring, complementing existing programs that generate long-term, management-relevant biodiversity data for MPA monitoring.

In this study, we assess the utility of a community science-based eDNA program for monitoring nearshore fish communities in two Southern California MPAs-the Point Dume State Marine Reserve and the Point Dume State Marine Conservation Area. Using two years of monthly sampling from volunteers associated with a local nonprofit Heal the Bay, we evaluate whether eDNA and community scientists can provide reliable, repeatable, and management-relevant biodiversity data. We specifically examine how sampling frequency, replication, and detection probability influence biodiversity inference for managers, and assess whether the two MPAs established in 2012 support different fish communities relative to nearby unprotected areas.

## Materials and Methods

### 2.1 Sample Collection

We conducted this study off the coast of Malibu, CA, under the State of California’s Natural Resources Agency Department of Fish and Wildlife scientific collection permit (SCP) number 13898. Sampling sites included the Point Dume State Marine Reserve (34.0024, -118.805) and its unprotected comparison site Latigo (34.0255, -118.756), and the Point Dume State Marine Conservation Area (34.0351, -118.8607) and its unprotected comparison site Piedra (34.0383, - 118.884) (Fig 1).

**Figure 1.**
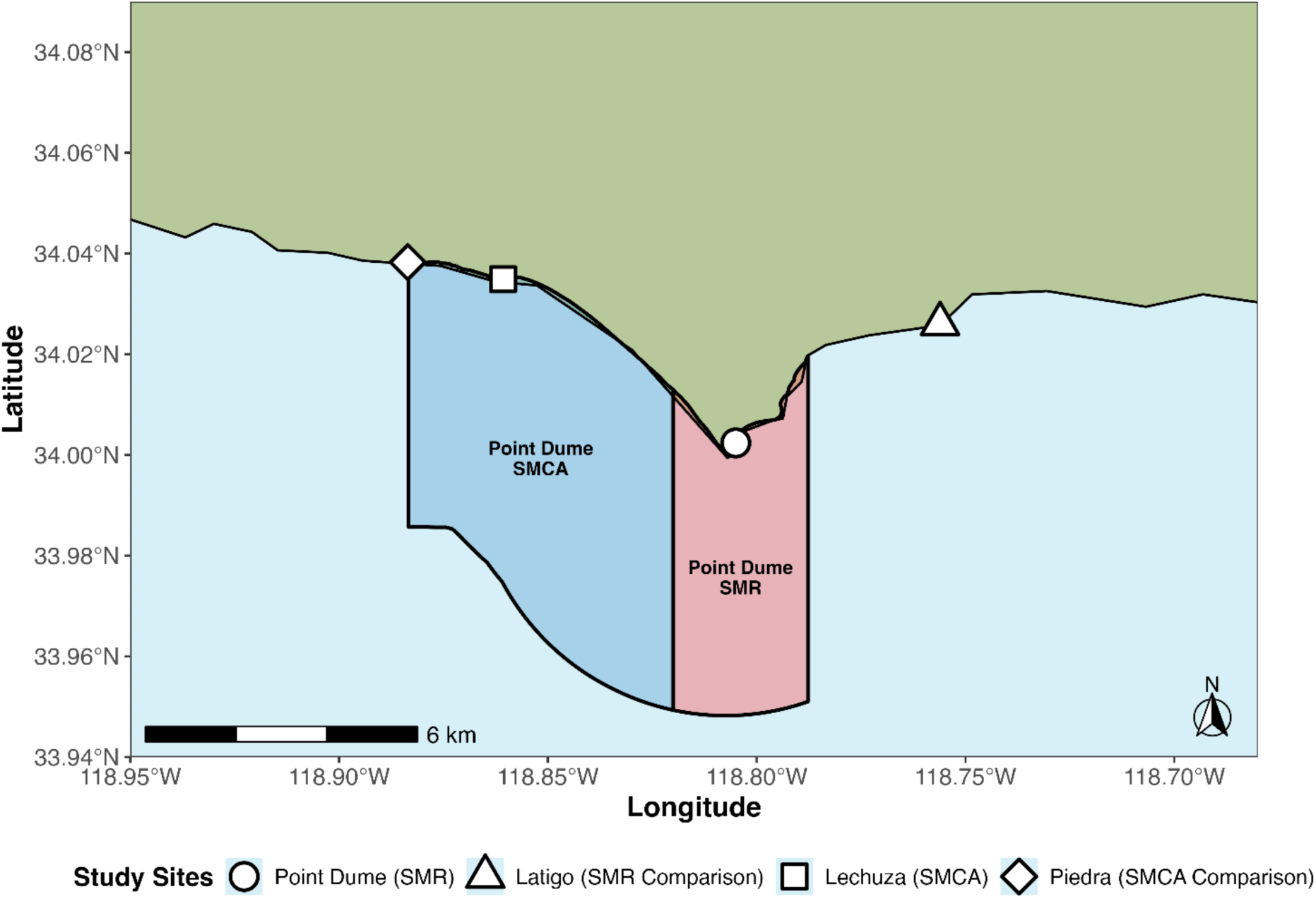
Map of the four sampling sites. Marine protected area boundaries are outlined. The map was generated using the free and open-source software R with packages sf and ggplot2.

Scientists, alongside our community nonprofit partner Heal the Bay, held quarterly training sessions for volunteers on MPA science, visual surveying techniques, and eDNA methodologies. Community scientists collected samples from April 2019 to March 2020. One month of data, April 2020, is missing from the dataset due to COVID-19 shutting down the beaches in LA County and making our sites inaccessible. From May 2020-April 2021, researchers alone took the monthly samples as COVID-19 guidelines limited field work by the community scientists.

We employed the eDNA collection methods outlined in Curd et al. 2019. The first Saturday of each month UCLA researchers met at a location central to the four sites and set up a congregation and filtration center. Heal the Bay volunteers arrived at 9am and were given a refresher training of how to collect environmental DNA samples. We organized volunteers into pairs, providing each with eDNA collection supplies (gloves, enteral feeding bags for water collection, and labelling markers) as well as a cooler and ice packs. Community scientists and researchers collected sweater samples at knee-depth off the beach at each site. We collected triplicate one-liter seawater samples in enteral feeding bags and kept water samples on ice to limit eDNA degradation. Within 20-30 minutes, scientists and volunteers returned to a central location where we filtered water samples through a sterile 0.22 µm Sterivex cartridge filter (MilliporeSigma, Burlington, MA, USA) using gravity filtration. We capped the filters and stored them on ice until we returned to the lab where they were stored at -20°C. For each sampling event, we filtered one liter of Milli-Q water through the same process for a negative field control.

### 2.2 DNA extraction and library preparation

We extracted eDNA from the Sterivex cartridges using a modified DNeasy Blood & Tissue Kit protocol (Qiagen Inc., Germantown, MD) optimized for increased eDNA yield^25^. Using PCR, we then amplified the extracted DNA using the Mifish Universal Teleost 12s primer set^26^. PCR reaction volume was 25 μL reaction composed of 12.5 μL QIAGEN Multiplex Taq PCR 2x Master Mix,1.5 μL of molecular grade H2O, 5 μL of forward primer (2 mM), 5 μL of reverse primer (2 mM), and 1μL of sample DNA. PCR cycling parameters employed a touchdown PCR profile with an initial denaturation at 95°C for 15 min, followed by 13 cycles of 95°C for 30s, beginning annealing at 69.5°C for 30 seconds and decreases in temperature of 1.5°C per cycle until 50°C, extension at 72°C for 1 minute. Initial touchdown cycles were followed by 24 cycles of 95°C for 30s followed by annealing at 50°C and an extension of 72°C for 1 minute. We use a final extension for 10 min at 72°C. All PCRs included a negative PCR control substituting molecular grade water for eDNA template. We verified PCR amplification success by checking product size on a 2% agarose gel electrophoresis stained with SybrSafe.

To create sequencing libraries, we indexed all successful PCR reactions using Nextera unique dual indices (Illumina, San Diego, CA, USA). Indexing PCRs used a 25 μL reaction composed of 12.5 μL Kapa Hifi MasterMix (Kapa Biosystems, Sigma Aldrich, St. Louis, MO, USA), 6.25 μL of molecular grade H_2_O, 1.25 μL of index and 5 μL of DNA from the PCR sample. Indexing PCR cycling parameters consisted of denaturation at 95°C for 5 min, followed by 8 cycles of denaturation at 98°C for 20s, annealing at 56°C for 30s, and extension at 72°C for 3 min, and then a final extension at 72°C for 5 min. We verified amplification success by checking product size on a 2% agarose gel electrophoresis stained with SybrSafe.

To ensure equal coverage in the sequencing libraries, we measured DNA concentration of each sample with the Qubit dsDNA Broad Range DNA Quantification Assay (Thermofisher Scientific, Waltham, MA, USA). We then pooled samples in equal molar concentrations in two sequencing libraries. To minimize artifacts related to batch effects, for each sampling period, sample replicate 1 was put on Run 1, sample replicate 2 was put on Run 2 and sample replicate 3 was randomized among Run 1 and Run 2. We cleaned the resulting library using a Qiagen gel extraction kit to remove both small and large non-target DNA fragments and sequenced the final library at the UC Davis’ Genome Center on an Illumina NovaSeq X (10B) using 150bp paired-end sequencing (PE150) spiked with 15% PhiX.

### 2.3 Bioinformatics

We used the *Anacapa Toolkit* for amplicon sequence variant parsing, taxonomic assignment, and quality control. Sequence processing began with trimming adapter sequences, removing low-quality reads, and sorting reads. DADA2 then parsed and dereplicated amplicon sequence variants (ASVs) prior to assigning taxonomy to using Bowtie 2 and a Bowtie 2-specific Bayesian Least Common Ancestor (BLCA) algorithm. Taxonomic assignment followed benchmarking parameters from Gold et al. (2021), using a taxonomic cutoff score of 60 and minimum alignment of 80%. Taxonomy was first assigned using the curated regional database of California Current Large Marine Ecosystem fishes to identify native taxa^27^. We then re-assigned the taxonomy using the global *CRUX* generated database to identify non-native and non-fish species. Taxonomic assignments of ASVs were synonymized between both methods by prioritizing higher resolution assignments (i.e. species level vs. genus level). In the samples that detected rockfish, we include “*Sebastes spp.”* because MiFish primers can’t resolve species within the genus *Sebastes* to species^28^. .

To eliminate poorly sequenced samples and remove potential sources of contamination, we implemented a decontamination procedure, employing a site occupancy modeling framework to retain only ASVs that occurred in high prevalence across locations and stations.

To provide insights into species relative abundance, we converted read counts into an eDNA index. The eDNA index transformation first normalizes all reads for a particular species by the total number of reads in each sample, then normalizes those proportions to the largest observed proportion for that species across all samples. This process results in a species-specific scaling between 0 to 1, where 1 is the sample with the highest occurrence for a given species in all the samples and 0 is the least. The index assumes that biases originate from DNA-primer interactions, which would remain consistent across samples, and thus allow us to infer relative abundance changes of a single taxa between samples^29^.

### 2.4 eDNA data analysis

To determine whether sequencing depth was sufficient to capture species richness in each sample, we generated rarefaction curves. From the rarefaction curves, we estimate the median number of reads needed for samples at each site to reach 95% of observed richness. To determine whether our field sampling captures site-level biodiversity, we used the iNEXT package^30^ to construct site-specific incidence-based species accumulation curves to estimate the minimum number of total samples needed to reach 95% of asymptotic species richness.

To evaluate how sampling effort could be reduced while retaining most of the biodiversity information captured by our intensive monthly sampling design, we used resampling simulations to compare alternative sampling frequencies. Using the complete two-year dataset, we simulate monthly, bi-monthly, and quarterly sampling strategies, as well as different combinations of seasons. For each strategy, samples were grouped by site, year, and the relevant sampling interval (i.e., monthly, two-month period, season), and we randomly selected one, two, or three samples from each interval. For each simulated dataset, selected samples were pooled and observed species richness was recalculated as the total number of taxa detected across selected samples. Each combination of sampling frequency and replication was resampled 500 times to account for variation within the samples and we summarized the mean species richness across iterations. We then compare these simulated richness estimates with the total richness observed under the full monthly sampling effort to determine how much sampling frequency and replication could be reduced to still maintain 95% species detection.

To estimate species detection probabilities, we used single-season occupancy models as implemented in the unmarked package^31^, assuming constant occupancy and detection. We constructed detection histories from replicate samples collected within the same site and sampling date. From these models, we calculated species-specific detection probabilities and the probability of detecting a species given one, two, and three replicate samples. To evaluate the relevance of eDNA detections for MPA management, we identified fish species of management concern based on the California Marine Protected Area Monitoring Action Plan for the South Region^32^. Species presence across sites and sampling periods was visualized using a heatmap by site and date to illustrate temporal and spatial patterns in species occurrence.

To compare eDNA detections with conventional visual surveys, we matched our eDNA samples with Reef Check California’s visual scuba surveys conducted at the same sites during the same month and year, although the surveys were not conducted concurrently or on the same day. Reef Check surveys target a pre-defined set of focal fish taxa rather than the full fish community. We therefore used two complementary comparisons. First, to evaluate how reliably eDNA recovered taxa independently observed by divers, we calculated the proportion of Reef Check-observed taxa that were also detected by eDNA for each matched site-month-year comparison. Because the MiFish 12s marker does not reliably resolve species within the genus *Sebastes*, all *Sebastes* detections were excluded from this analysis. Second, to compare the breadth and type of taxonomic information provided by each method, we retained all detections, including *Sebastes*, and classified taxa within each matched comparison as either detected by both methods, detected only by Reef Check, or detected only by eDNA.

To evaluate the impacts of MPA status on fish populations, we calculate alpha diversity as total observed species richness for each sample and compared richness between each MPA and its paired comparison site. We conducted analyses separately from the State Marine Reserve (SMR) and the State Marine Conservation Area (SMCA), comparing total species richness using a negative-binomial generalized linear model with site (i.e MPA or non-MPA), season, and year as predictors. The significance of each predictor was evaluated using likelihood-ratio tests, and model fit was assessed using Pearson residual dispersion. To evaluate relative abundances in each MPA and its comparison site, we compared median species-specific eDNA index values using a nonparametric Mann-Whitney U test. To account for multiple testing across species, we adjusted p-values to correct for false discovery rates.

To assess associations between species richness and environmental variability, we evaluated five environmental variables’ impacts on observed species richness: the Multivariate El Niño Southern Oscillation Index (ENSO index^33^), Pacific Decadal Oscillation (PDO^34^), the Biologically Effective Upwelling Transport Index (BEUTI^35^), the Coastal Upwelling Transport Index (CUTI^35^), and peak sea surface temperature (SST^36^). Each variable was first evaluated using separate linear regressions of monthly richness. We then fit a focused linear model testing the relationship between richness and the highest correlated variable while controlling for site and season.

To test for differences in community composition (beta diversity), we converted eDNA indexes for all samples into Bray-Curtis dissimilarity distances from eDNA index values. We then tested for differences in community structure by site, season, and year using a PERMANOVA and assessed homogeneity of multivariate dispersions among sites using BETADISPER. Lastly, we visualized community composition using a constrained analysis of principal coordinates (CAP) based on Bray-Curtis distances, constraining ordinations by site and season.

To evaluate the feasibility of eDNA MPA monitoring in statewide implementation, we estimated the per-sample cost of each step in the eDNA workflow used in this study including field sampling, DNA extraction, PCR amplification, gel electrophoresis, sample indexing, bead cleaning, DNA quantification, and sample sequencing (Table S5). Original prices were based on costs at the time of sampling in 2019. Prices were adjusted to account for inflation to approximate present-day values although these remain estimates as not all costs increase the same as general inflation.

## Results

### 3.1 Species richness and sampling sufficiency

Sequencing on the NovaSeq X, combined with quality control in the Anacapa Toolkit, resulted in 1,231,029,727 reads. After taxonomic assignment, 844,768,875 reads remained across 249 samples, representing 63 families, 103 genera, and 119 species. Each sample’s ASV read counts and eDNA index values are reported in Supplementary Table 1 and 2.

Rarefaction curves showed that for each sample, sequencing depth was sufficient to capture all species diversity within that sample (S1 Fig). Site-specific rarefaction curves modeled using the iNext package show that samples at each site was sufficient to fully capture local diversity (S2 Fig), and indicate that approximately 11-14 samples are needed to reach 95% of estimated effective number of species at each site (Latigo (SMR Comparison): 11, Point Dume (SMR): 14, Piedra (SMCA Comparison): 11, Lechuza (SMCA): 13). Using sample-level rarefaction curves, we estimated the median number of reads required for samples at each site to reach 95% of observed richness to be between 705k-990k reads (Latigo (SMR Comparison): 990k , Point Dume (SMR): 762k, Piedra (SMCA Comparison): 705k, Lechuza (SMCA): 775k).

Resampling-based simulations showed that species richness increased with both sampling frequency and the number of replicate samples taken (Table 1). Monthly sampling across the full duration of the study recovered the highest mean observed species richness. Moreover, richness increased from 112.2 species with one replicate to 117.5 species with two replicates, and peaked at 119 species with three replicates (94.3%, 98.7%, and 100% of observed richness, respectively). Bi-monthly sampling recovered less total diversity, although diversity still increased with each additional replicate from 105.9 to 112.6 to 116.2 species (89%, 94.6%, and 97.6% of observed richness, respectively). This pattern was consistent with quarterly sampling, with richness increasing from 104.2, to 111 to 114.4 species with each additional replicate (87.6%, 93.3%, and 96.2% of observed richness, respectively). Seasonal sampling regimes recovered fewer species overall but up to 95% of total diversity found sampling monthly. Sampling across three seasons with three replicates recovered between ∼111-113 species (93-95% of observed richness), two season sampling recovered ∼106-111 species (89-93% of observed richness), and single season sampling recovered between ∼97-102 species (81-86% of observed richness).

**Table 1:** Effects of sampling frequency and bottle replication on estimated species richness based on resampling simulations of our March 2019-April 2021 sampling. Sample counts for each strategy were generated by binning observations at the site x season x year level and drawing an equal number of samples per bin. Because the study period spans March 2019-April 2021, some seasons occur in more calendar years than others (e.g. three winters and springs but only two summers and falls), resulting in different total sample counts among single-season strategies.

| Sampling Strategy | Number of<br>Samples -<br>1 Replicate | Number of<br>Samples -<br>2 Replicates | Number of<br>Samples -<br>3 Replicates | Mean<br>Richness-1 Replicate | Mean<br>Richness-2 Replicates | Mean Richness-3 Replicates |
| --- | --- | --- | --- | --- | --- | --- |
| Monthly | 92 | 184 | 249 | 112.208 | 117.496 | 119 |
| Bi-monthly | 48 | 96 | 144 | 105.854 | 112.62 | 116.194 |
| Quarterly | 40 | 80 | 118 | 104.248 | 111.012 | 114.29 |
| Winter + Spring + Summer | 40 | 80 | 118 | 104.226 | 111.014 | 114.434 |
| Winter + Spring + Fall | 32 | 64 | 94 | 102.464 | 109.304 | 112.938 |
| Spring + Spring + Fall | 32 | 64 | 94 | 101.092 | 107.468 | 110.858 |
| Winter + Spring + Fall | 28 | 56 | 82 | 100.7 | 106.86 | 110.794 |
| Winter + Spring | 28 | 56 | 84 | 100.576 | 107.698 | 111.822 |
| Winter + Summer | 24 | 48 | 70 | 98.294 | 104.81 | 108.598 |
| Spring + Summer | 20 | 40 | 60 | 98.184 | 105.436 | 109.726 |
| Spring + Fall | 20 | 40 | 58 | 98.022 | 104.546 | 107.99 |
| Spring + Fall | 20 | 40 | 58 | 96.096 | 102.44 | 105.37 |
| Winter + Fall | 20 | 40 | 60 | 95.938 | 102.444 | 106.164 |
| Summer + Fall | 16 | 32 | 48 | 95.008 | 102.176 | 106.47 |
| Winter | 12 | 24 | 36 | 90.462 | 97.734 | 102.022 |
| Spring | 12 | 24 | 34 | 89.868 | 96.624 | 99.286 |
| Summer | 8 | 16 | 24 | 88.568 | 96.862 | 101.396 |
| Fall | 8 | 16 | 24 | 84.866 | 92.902 | 96.552 |

### 3.2 Species detection

Species occupancy models were fit for 100/119 taxa and estimated species-specific occupancy and detection probability. Occupancy probabilities ranged from 0.0992-1 (median 0.9928) with 62 taxa exhibiting occupancy greater than or equal to 0.95. Single-replicate detection probabilities for species ranged from 0.0202-1 (median 0.4175) (Fig 2; Table S4). Detection probabilities increased with each eDNA sample replicate, with a median of 0.6604 (range: 0.0400-1) for two replicates and a median of 0.8018 (range 0.0595-1) for three replicates. Across taxa, detection probabilities with three replicates exceeded 0.95 for 32 species, fell between 0.5 and 0.95 for 51 species, and fell below 0.5 for 17 species (Table S4).

**Figure 2.**
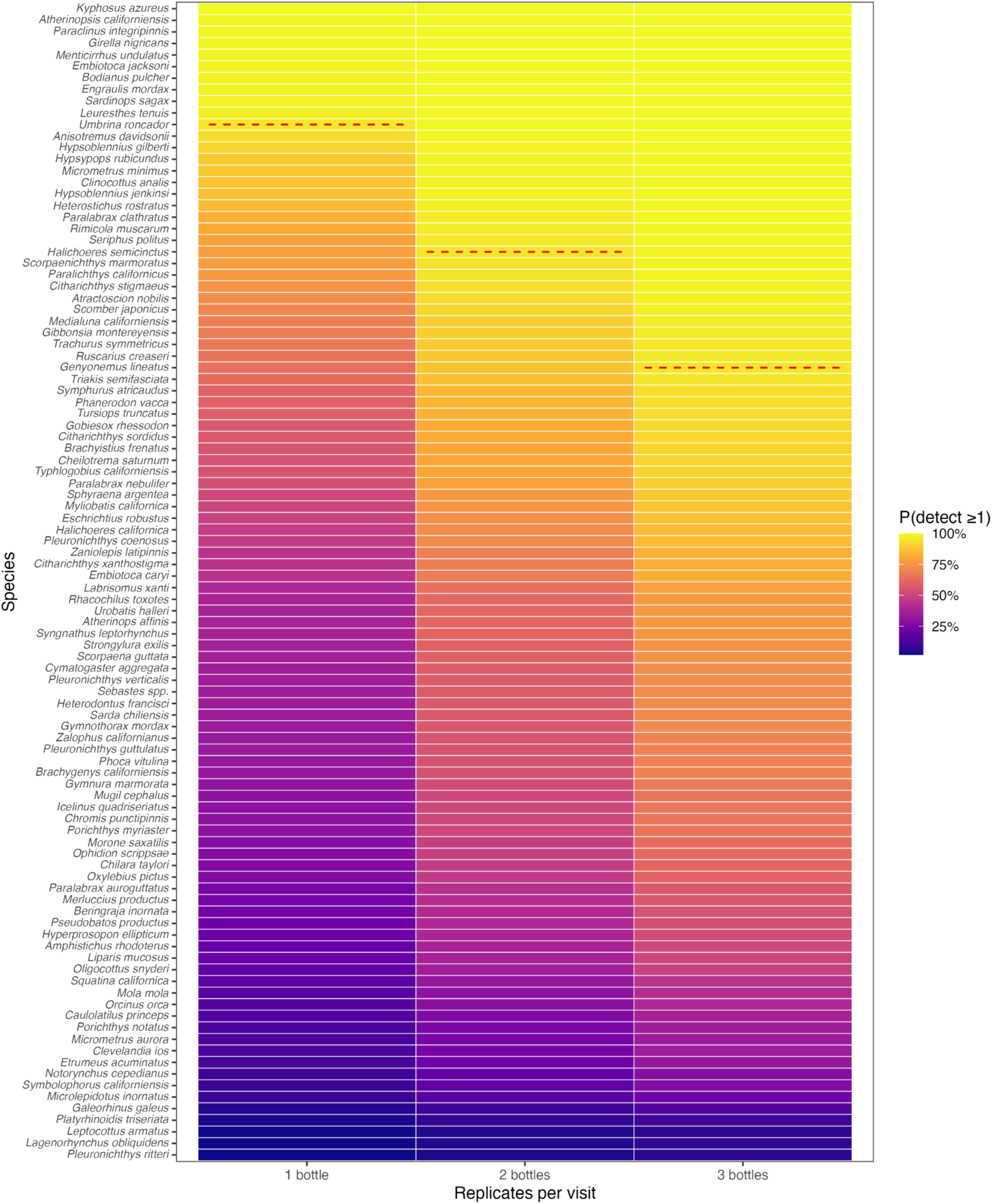
Estimated detection probabilities across 100 taxa based on species occupancy models with 95% detection probability indicated using a red dashed line, showing the probability of species detection increases with the number of sampling replicates..

Across all sites and sampling days, eDNA detected 19/22 non-rockfish species identified as species of management interest in the California Marine Protected Area Monitoring Action Plan for the Southern Region (S3 Table), only failing to detect *Paralabrax maculatofasciatus, Ophiodon elongatus,* and *Anoplopoma fimbria* which would not be expected in these nearshore, open coast sites. In addition to these 22 species of management interest, our eDNA marker also detected 8 shark species (*Alopias vulpinus*, *Cephaloscyllium ventriosum*, *Cetorhinus maximus*, *Galeorhinus galeus*, *Heterodontus francisci, Mustelus californicus*, *Notorynchus cepedianus* and *Triakis semifasciata*), 5 ray species (*Beringraja inornata*, *Gymnura marmorata*, *Myliobatis californica*, *Pteroplatytrygon violacea*, *Urobatis halleri*) and 8 marine mammals species (*Eschrichtius robustus*, *Lagenorhynchus obliquidens*, *Megaptera novaeangliae*, *Mirounga angustirostris*, *Orcinus orca*, *Phoca vitulina*, *Tursiops truncatus*, *Zalophus californianus*).

### 3.3 MPA vs non-MPA comparisons

Analyses showed lower observed species at both MPA sites relative to the non-MPA sites (Fig 3). In the SMR pair, mean observed richness was 42.4 species at the MPA (Point Dume) and 48.3 species at the comparison site (Latigo), a mean difference of 5.9 species. After accounting for season and year, expected richness at Point Dume was 12.5% lower than at Latigo (ratio rate= 0.975, 95% CI= 0.766-1.001), although the site effect was not statistically significant (likelihood-ratio test, p=0.052). In the SMCA pair, mean observed richness was 43.5 species at the MPA (Lechuza) and 48.5 at the comparison site (Piedra), a mean difference of 5 species. Expected richness at Lechuza was 11.5% lower than at Piedra after accounting for season and year (rate ratio= 0.885, 95% CI= 0.785-0.998), and the site effect was statistically significant (likelihood-ratio test, p = 0.048). Season was associated with richness in the SMCA pair (p= 0.020) but not the SMR pair (p=0.931) while year was not significant in either model (p= 0.414 and p= 0.305, respectively).

**Figure 3:**
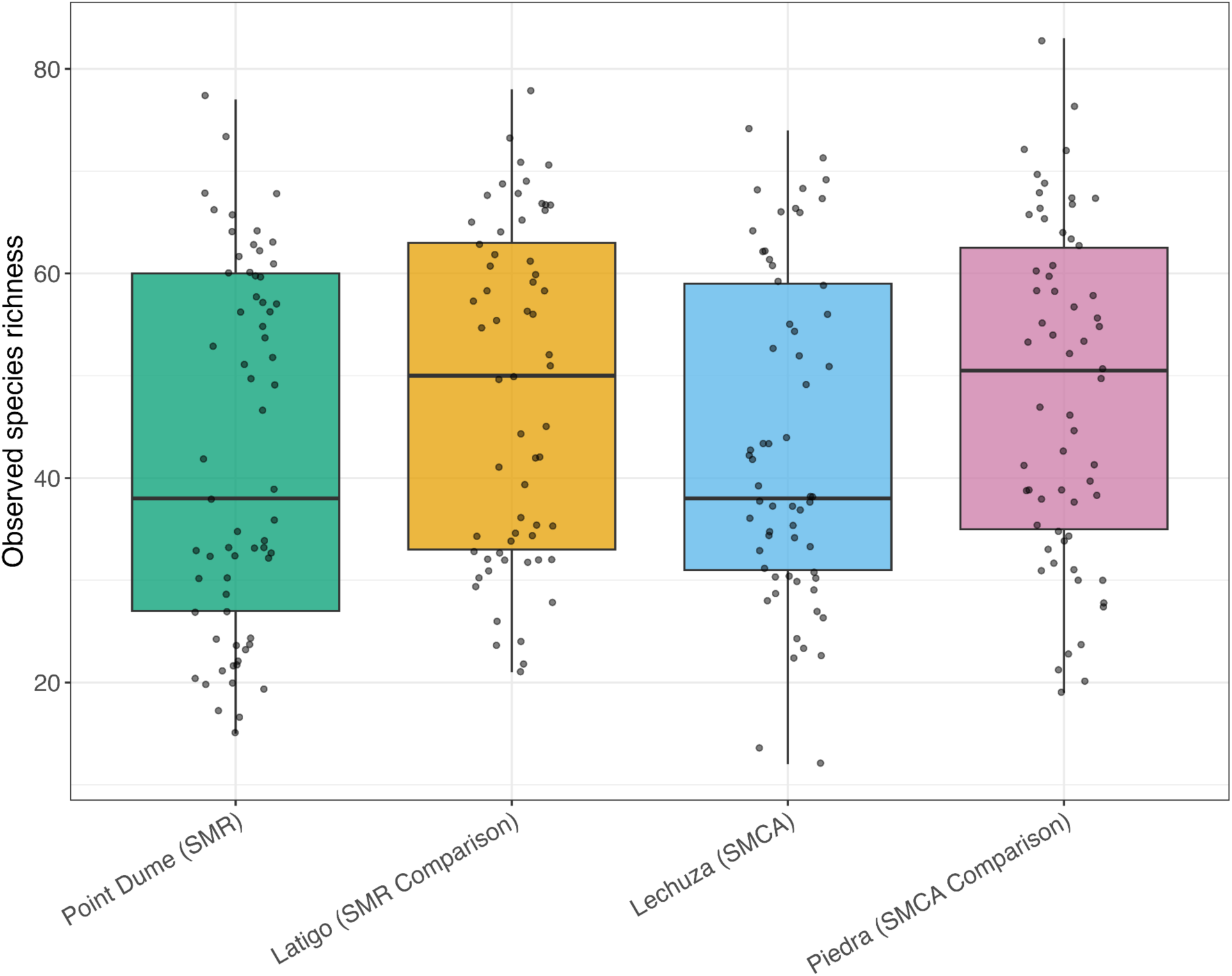
Observed species richness across the four sampling sites. Both comparison sites had higher mean observed richness than their paired MPA sites. Negative-binomial generalized linear models accounting for season and year detected a significant site effect for the SMCA pair (p=0.048), while the SMR site effect was not statistically significant.

Although observed species richness trended higher at comparison sites than MPAs, species composition was highly similar across all sites. A total of 119 taxa were detected with 93 species (78.2%) detected at all four sites (Fig 4). Temporal presence-absence patterns further showed that each site was consistently characterized by the same set of common nearshore taxa across much or all of the sampling period (Fig 4). Taxa restricted to MPA sites were limited to six species including two unique to the SMCA (*Stereolepis gigas* and *Synodus lucioceps*) and four shared between the two MPAs but absent from their comparison sites (*Cephaloscyllium ventriosum*, *Lyposetta exilis*, *Syngnathus exilis*, and *Zalophus californianus*).

**Fig 4.**
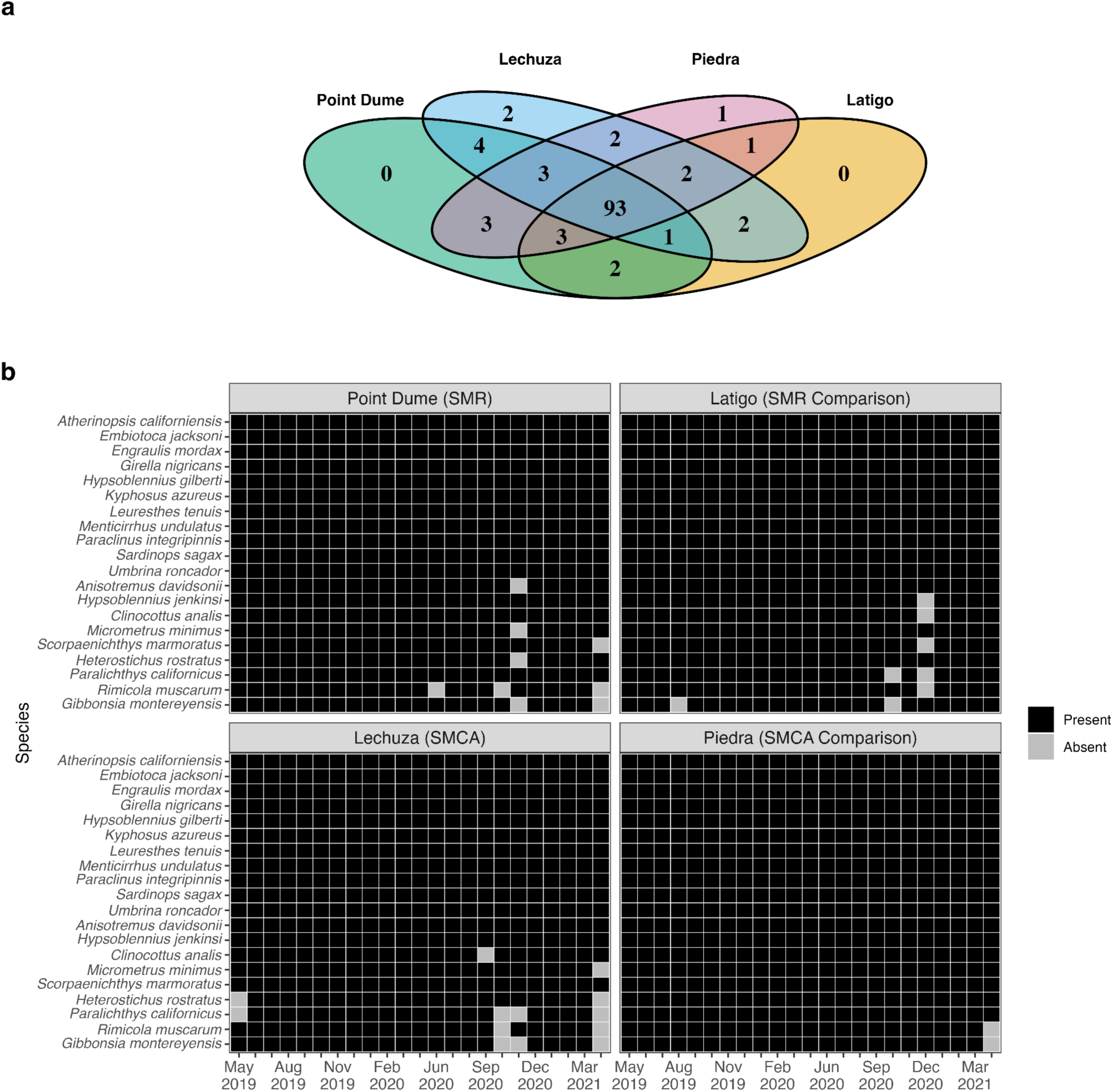
a) Four-site Venn diagram showing overlap in species detections among sites, with most taxa shared across all four sites and relatively few taxa unique to any individual site. b) Spatial and temporal patterns of species detections across sites and sampling days of the top 20 species highlights the similarities between sites.

### Time series analyses

Across the 24-month sampling period, monthly species richness was generally stable across all four sites with most samples ranging between 55-80 species (Fig 5). Richness patterns were broadly similar across sites and exhibit limited seasonal variation. The most pronounced change in this pattern occurred during the start of the 2020 La Niña cycle, which started August 2020 and was associated with richness declining to a low of 35-45 species at Latigo, Lechuza, and Point Dume before recovering to prior levels of around 70 species three months later (Fig 5).

**Fig 5.**
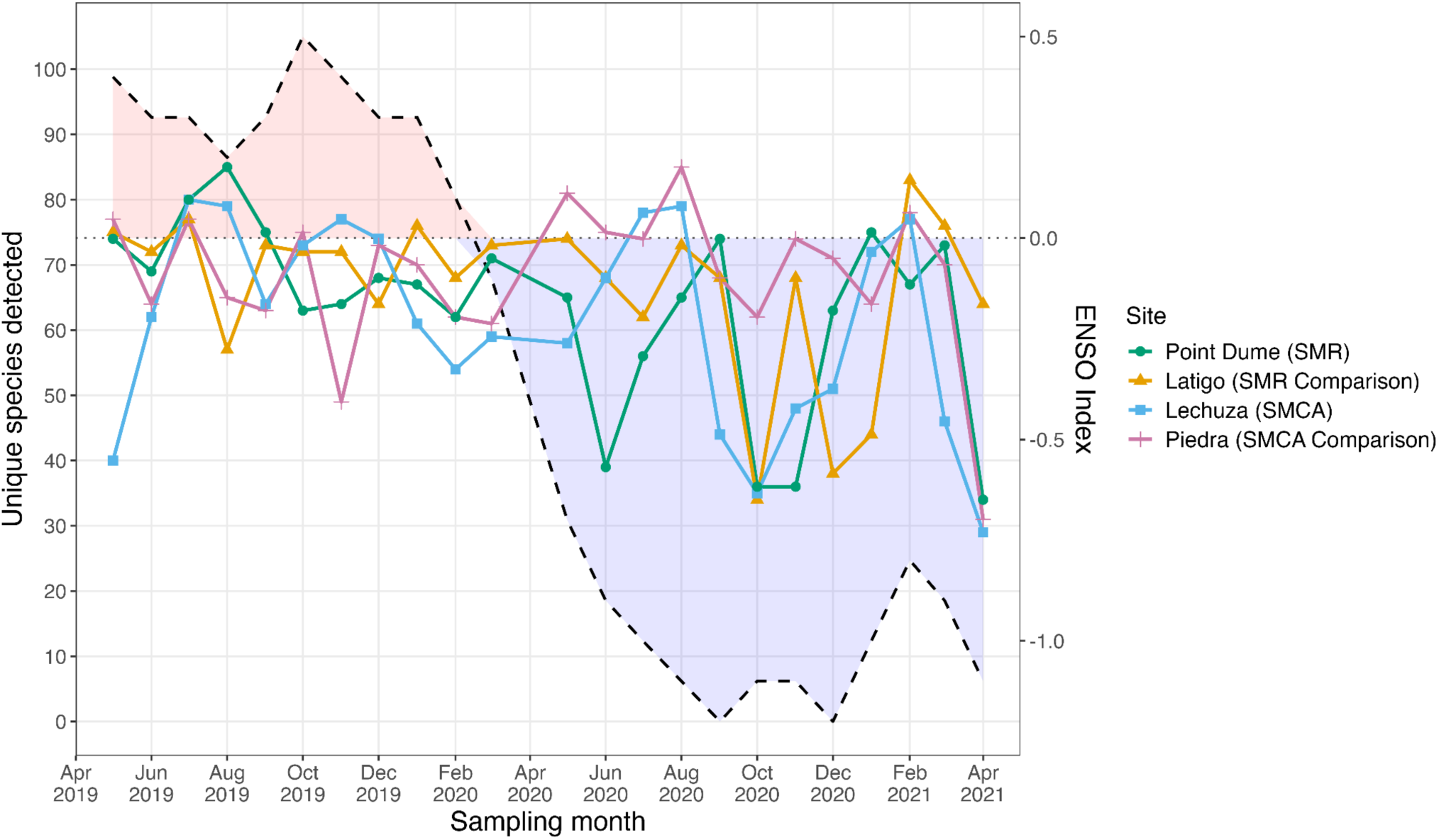
Monthly species richness across sampling sites with a dashed line indicating NOAA’s ENSO index. Species richness was generally stable prior to La Niña, declined across sites during the onset of negative ENSO conditions, and recovered in subsequent months. At Lechuza, community composition was significantly correlated with ENSO index in regression analyses (Figure S3).

To evaluate potential drivers of this temporal variability, monthly richness was modeled as a function of environmental variables while controlling for site and season. Among the variables tested, NOAA’s Multivariate El Nino Southern Oscillation Index MEI.V2 (ENSO index) was the strongest predictor (S3 Fig), with site and season explaining 11.9% of richness variation; inclusion of the ENSO index increased explained variance to 21.1% (p = 0.0023). In contrast, the Bakun upwelling index, CUTI, BEUTI, Pacific Decadal Oscillation, and peak SST were not significantly correlated to richness.

To assess whether the 2020 La Niña event was associated with changes in community composition, samples were grouped into pre-La Niña (May 2019-August 2020) and during La Niña (September 2020-April 2021). Samples at each site pre-La Niña and during La Niña were compared using CAP ordination (S4 Fig). When controlling for season within each site, seasonal effects explained between 6-9.7% of variation in community composition and were significant at Point Dume (R^2^ = 0.097, p < 0.001), Lechuza (R^2^ = 0.081, p < 0.001), and Piedra (R^2^ = 0.097, p < 0.001), but were not significant at Latigo (R^2^ = 0.060, p = 0.077). In contrast, the La Niña phase was not significantly associated with community composition at Point Dume (R^2^ = 0.008, p = 0.978), Latigo (R^2^ = 0.021, p = 0.125), or Piedra (R^2^ = 0.018, p = 0.175). Only Lechuza showed a modest difference between pre- and during-La Niña periods (R^2^ = 0.027, p =0.014). However, multivariate dispersion also differed significantly at Lechuza (Betadisper, p = 0.025), indicating that this result reflects greater within-period variability rather than directional shift in community composition.

Community composition differed between MPA and non-MPA sites for both pairs. For the SMR, Bray-Curtis dissimilarities varied by site, season, and year (PERMANOVA, R^2^ = 0.116, p= 0.001) with significant effects of site (R^2^ = 0.034, p= 0.001), season (R^2^ = 0.048, p= 0.002), and year (R^2^ = 0.030, p= 0.001). However, multivariate dispersion differed between the two sites (BETADISPER, F= 22.24, p < 0.001). CAP ordination constrained by site and season was significant overall with both terms contributing significantly (p=0.001). For the SMCA, Bray-Curtis dissimilarities also varied by site, season, and year (PERMANOVA, R^2^ = 0.099, p= 0.001) with significant effects of site (R^2^ = 0.025, p= 0.001), season (R^2^ = 0.044, p= 0.001), and year (R^2^ = 0.025, p= 0.002). Dispersion did not differ between the SMCA and its comparison (BETADISPER, p= 0.116). CAP ordination constrained by site and season was significant overall with both terms contributing significantly (p=0.001).

To evaluate whether rare taxa were driving differences in these patterns, we compared CAP ordination plots generated using the full set of species to plots generated based on the top 10 most abundant species found across sites. The two ordinations were highly concordant (Procrustes correlation = 0.74, p= 0.0001), indicating that observed site and seasonal differences were primarily driven by common species rather than sporadic detections of rare taxa.

Although statistical tests were conducted separately for each MPA-comparison pair to account for site differences, all four sites are visualized together in a single CAP ordination plot to facilitate comparison of community patterns across the study region. (Fig 6)

**Figure 6.**
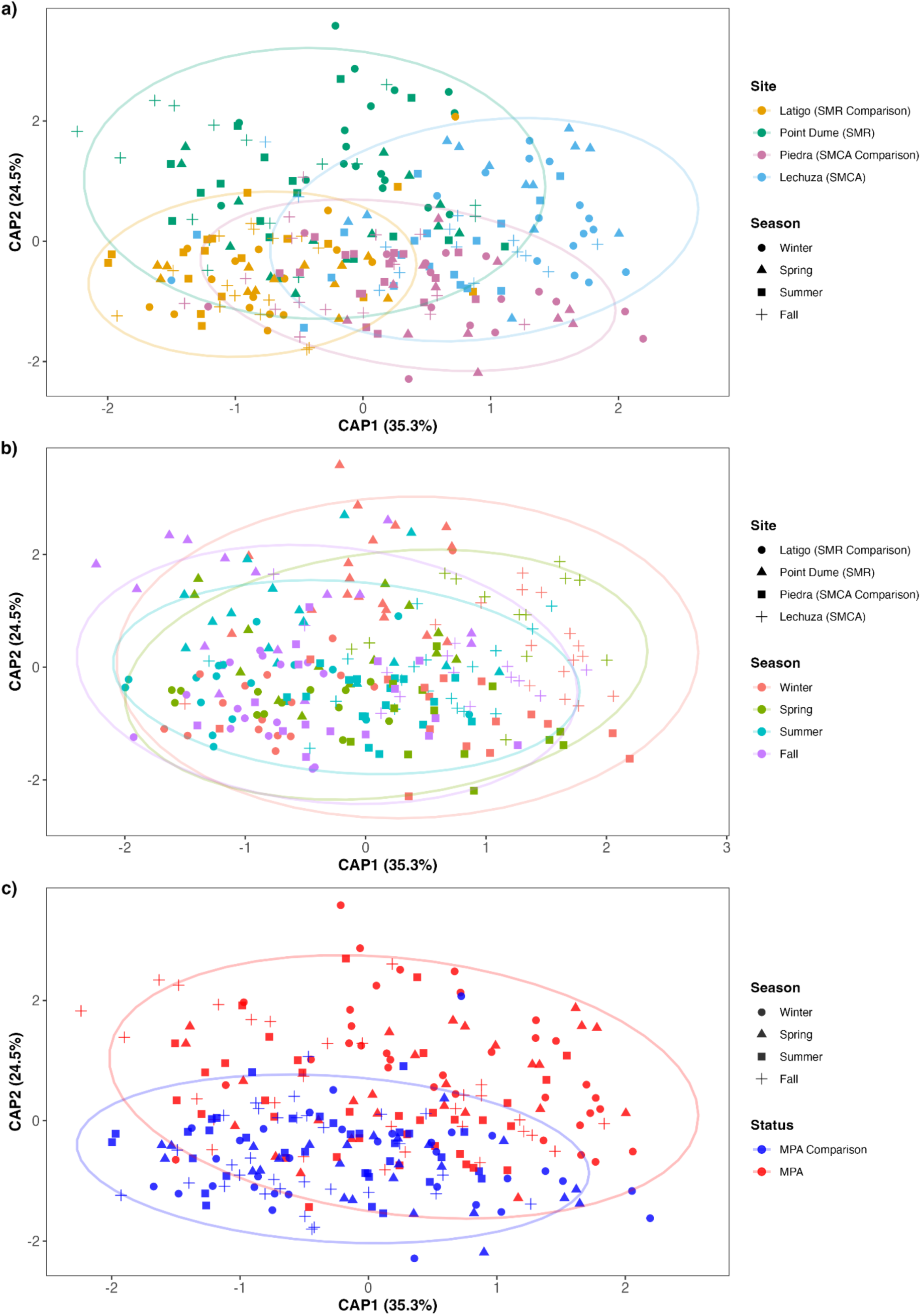
CAP ordination of community composition across four sites, colored by (a) site, (b) season, and (c) MPA status. Community structure clustered most strongly by site followed by season and then MPA status.

Only a small number of fish species showed statistically significant differences in relative abundance between MPA and the comparison site. At the SMR site pair, seven species showed significantly higher relative abundances. Of the seven, six species (*Heterostichus rostratus* (p<0.001), *Triakis semifasciata* (p<0.001), *Embiotoca jacksoni* (p=0.001), *Rimicola muscarum* (p<0.001), *Cheilotrema saturnum* (p<0.001), and *Paraclinus integripinnis* (p<0.001)) had higher relative abundances at the non-protected site and only one species (*Atractoscion nobilis* (p<0.001)) had higher relative abundance at the MPA site. At the SMCA site pair, six species showed differences in relative abundances. All six species (*Girella nigricans* (p<0.001), *Heterostichus rostratus* (p<0.001), *Paraclinus integripinnis* (p=.002), *Rimicola muscarum* (p=.02), *Triakis semifasciata* (p=.0046), and *Seriphus politus* (p=.0046)) showed higher relative abundance at the comparison site.

### 3.4 Comparison to scuba surveys

Six Reef Check surveys could be matched to eDNA sampling conducted at the same site within the same month and year. After excluding *Sebastes*, eDNA detected a mean of 83.5% of the taxa observed by Reef Check (median = 87.3%, SD= 13.5%, range = 61.5-100%). Across all six comparisons, only seven unique non-*Sebastes* taxa observed by Reef Check were not detected by eDNA. *Heterodontus francisci* was missed by eDNA in three matched surveys; *Chromis punctipinnis* and *Halichoeres californica* were each missed in two; and *hypsurus caryi, Medialuna californiensis, paralabrax nebulifer,* and *Rhacochilus toxotes* were each missed in one. When all taxa were retained for both methods, the two approaches differed substantially in the breadth of species richness detected (Fig. 7). Across each of the six matched comparisons, eDNA detected considerably more taxa that were not observed by Reef Check.

**Figure 7:**
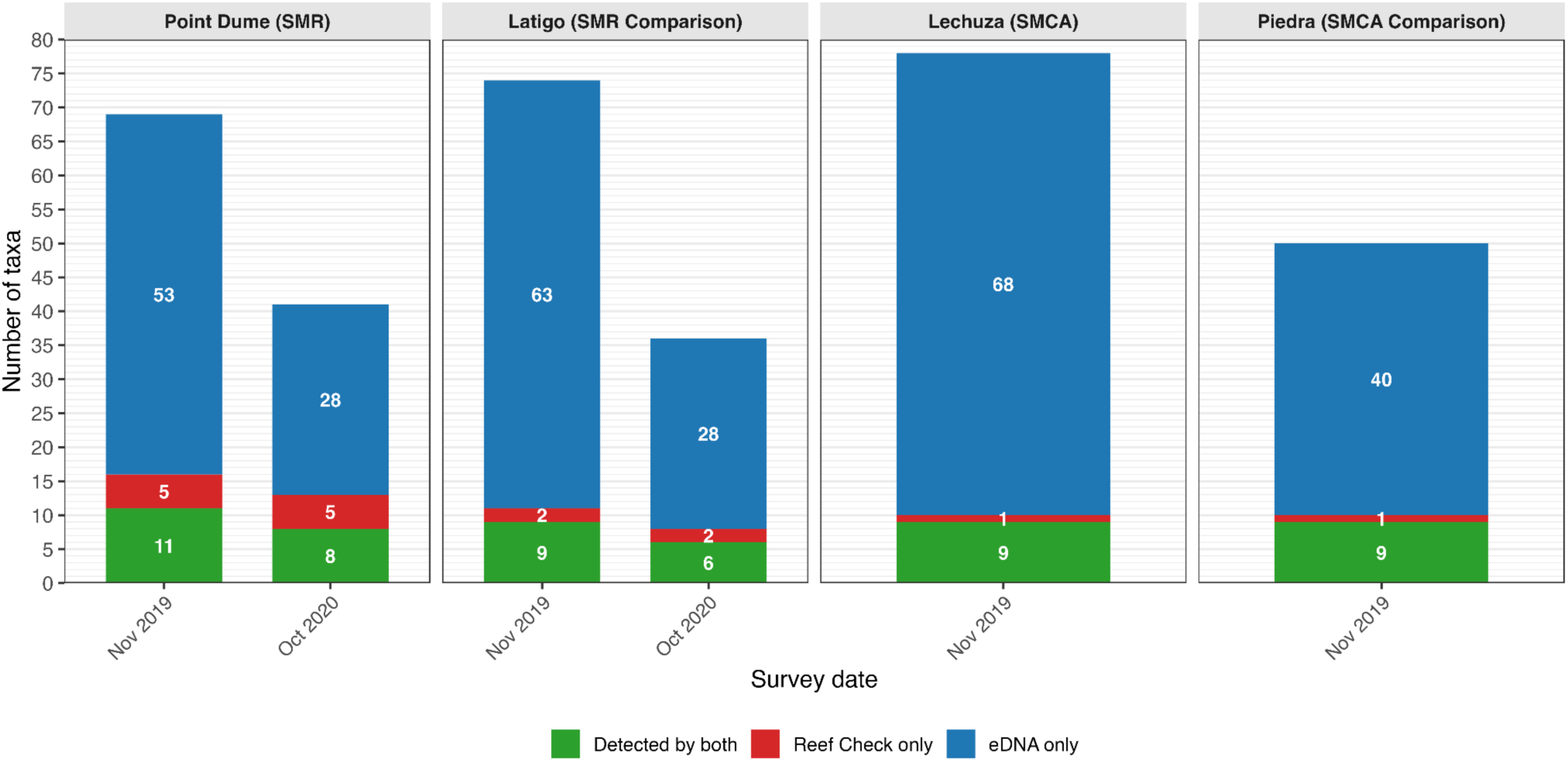
Comparison of taxonomic detections from eDNA metabarcoding and Reef Check visual surveys across six matched site-month-year sampling periods. Stacked bars show taxa detected by both methods, detected only by Reef Check, and only by eDNA. Numbers within each bar segment indicate the number of taxa in that category. Across all six comparisons, eDNA detected more taxa than were recorded by the Reef Check surveys.

### 3.5 Scaling eDNA to all 124 MPA sites

To assess the feasibility of scaling this eDNA workflow statewide, we estimated the present-day cost per sample of the full pipeline from sampling to sequencing. Across all stages, the estimated adjusted cost was roughly $52.38 per sample. We then scaled these costs to California’s 124 MPAs and paired reference sites using the minimum sampling design identified by this study: three replicate samples per site collected quarterly. This design would generate 1,488 MPA samples (124 sites x 3 replicates x 4 sampling events) and 1,488 comparison site samples. We additionally included one field blank per site pair during each quarterly sampling event, increasing the total number of samples to 3,472 annually. Because field blanks would only proceed to indexing and sequencing if contamination was detected, downstream library prep was calculated using the 2,976 biological samples rather than all samples. Under these assumptions, annual supply costs were estimated at approximately $172,000. To account for sample processing, data management, and analysis, we also include the annual salary of two full-time State Environmental Scientist (A) at $68,975 each. Combined, these costs yield an estimated annual baseline cost of roughly $310,000. This estimate assumes that field collections are integrated into existing community science networks across the state and therefore does not include dedicated labor costs for sample collection.

## Discussion

Our results demonstrate that community science-based eDNA sampling can provide scalable, management-relevant biodiversity information for long-term monitoring of California’s marine protected areas. Across two years of sampling, eDNA detected 119 species, including 19 or 22 regional management-priority fish species with community scientist collected samples producing biodiversity data comparable to that of samples collected by researchers. eDNA also recovered most taxa that were independently observed in matched scuba surveys while detecting a substantially broader set of taxa overall. Our analyses further showed that much of the biodiversity information obtained through intensive monthly monitoring could be retained with substantially lower sampling effort and sufficient sequencing depth. Together, these findings support eDNA as a practical complement to established visual monitoring programs with the potential to expand both spatial and temporal coverage of California’s MPA monitoring network at low-costs.

### 4.1 eDNA monitoring reliably detects management-relevant taxa across Southern California MPAs

This study demonstrates that volunteer-collected eDNA samples can produce high-quality, management-relevant biodiversity data. Our eDNA surveys were able to detect 119 species across the four sites with 109 unique species found at Point Dume State Marine Reserve and 109 unique species found at Point Dume State Marine Conservation Area. From these surveys, our samples detected 19 of the 22 non-rockfish species designated as management priorities in the Southern Region MPA Monitoring Action Plan across the four sites. The three undetected species, *Paralabrax maculatofasciatus*, *Ophiodon elongatus*, and *Anoplopoma fimbria* , would not be expected to be captured in nearshore sampling at Point Dume as it falls outside their respective range or habitat type.

Additionally, our eDNA surveys were able to detect other species with significant management concern. This includes the IUCN critically endangered giant black seabass (*Stereolepsi gigas*) and the IUCN endangered basking shark (*Cetorhinus maximus*). It also includes both commercially and recreationally important fish to California. For commercial species, our surveys detected California halibut (*Paralichthys californicus*), the 5th largest fishery by ex-vessel value at $5.7 million, and the northern anchovy (*Engraulis mordax*), Pacific sardine (*Sardinops sagax*), and Pacific mackerel (*Scomber japonicus*), the 3rd, 4th, and 5th largest fishery by weight^37^. For recreational species, eDNA detected almost all of the top 10 species of fish targeted by recreational anglers (based on pounds of fish caught) with the exception of numbers 1 (Tunas) and 8 (Yellowtail)^37^. Beyond this, while our study focused primarily on teleost fishes, our eDNA surveys were also able to detect 8 shark species, 5 ray species, and 8 marine mammals. Environmental DNA was able to track the presence and absence of these priority species across sites and months, providing spatially and temporally resolved information that can help inform management decisions and help managers better understand patterns of occurrence. We also highlight that future work could expand the number of species detected by employing a wider range of metabarcoding loci. For example, additional loci could be added to target rockfishes^28^, decapods^38^, and broad metazoan diversity^39^.

For a natural resource agency, detection reliability is an equally important consideration for eDNA’s integration into management use. With the statistical power of a large dataset across two years of monthly sampling, we were able to create a robust profile of detection probabilities for the species detected at our study sites. The species in this study had a median probability of detection of 0.802 when three replicates were taken. 32 species had a 95% detection probability, 51 additional species had a probability between 50%-95%, and 17 species had less than a 50% detection probability. The ten fish with the lowest detection probability consisted of deep-water pelagic fish (*Symbolophorus californiensis*), which would be difficult to capture from the nearshore sampling setup this study used, estuary associated fish species (*Clevelandia ios*, *Leptocottus armatus*), which are less common in an open-coast reef environment, and sharks, rays, and marine mammals (*Galeorhinus galeus*, *Lagenorhynchus obliquidens*, *Notorynchus cepedianus, Platyrhinoidis triseriata*), which have large home ranges, are more transient, and have a better amplification efficiencies and resolution using more targeted primer set, such as the MiFish-elasmobranch primer^26^.

Beyond documenting species of management interest, repeated eDNA sampling provides managers with the ability to evaluate how nearshore communities are responding to changing ocean conditions. During the 2020-2021 La Niña transition, eDNA documented declines in species richness at three of the four sites. Although the underlying mechanisms for this pattern cannot be resolved from our dataset, La Niña associated conditions influence fish community composition and distribution through several non-mutually exclusive pathways. First, cooler ocean temperatures can cause shifts in distribution of temperature-sensitive species, as observed in flatfish along the U.S. West Coast during periods of cooler water^40^. Second, because eDNA shedding rates are linked to water temperature, colder conditions may decrease eDNA shedding rates and lower detection rates for low-abundance species^41^. Third, stronger upwelling and changes in circulation during La Niña conditions may increase water column mixing and reduce the retention of eDNA from the sampling sites, which would dilute site-specific signals. The ability to track changes in fish community assemblages over space and time is important for understanding climate range shifts, tracking invasives, monitoring species of conservation concern, and providing baseline data to understand impacts from episodic events, such as major wildfires (e.g. the 2025 Palisades fire) or the anticipated 2026-27 ‘super’ El Niño.

### 4.2 Incorporating community scientists expands the spatial and temporal coverage of biodiversity monitoring while reducing time and effort

Environmental DNA metabarcoding proved highly effective at detecting fish assemblages across both the protected and unprotected sites in this study. The performance of eDNA was consistent during sampling periods led by community sciences as well as those conducted by university researchers during Covid-19 when our volunteer program was temporarily paused. During the first year of the project, approximately 70 community scientists contributed to collecting 144 eDNA samples, substantially increasing sampling capacity relative to what would have been feasible through a research team alone with the same budget. Through the participation of community members in biodiversity monitoring, we expanded both the scope and scale of sampling while reducing the amount of time and effort on researchers.

This expanded capacity is particularly important in the context of statewide habitat monitoring, where conventional survey approaches can be prohibitive to scale. California’s MPA network encompasses 124 sites distributed across geographically and biologically diverse coastline. Conventional approaches for monitoring fish assemblages include scuba-based visual surveys, baited remote underwater videos (BRUV), and net sampling. While these methods provide important information on traits known to be affected by MPA status, such as abundance, biomass, and size, they require specialized knowledge and equipment making them difficult to quickly and easily implement statewide.

Existing programs effectively harness the power of community science, such as Reef Check (www.reefcheck.org), a community science-based scuba survey program that trains and organizes volunteer divers to collect biodiversity information about reefs. However, there is a barrier to participation in scuba-based surveys as it requires a costly scuba diving certification as well as specialized taxonomic knowledge of local species for proper identification^42,43^. While programs like Reef Check are critical for monitoring our oceans, these requirements restrict the potential pool of participants and limit the scale at which community-based biodiversity monitoring can occur. Additionally, scuba surveys are time- and energy-intensive in the field, restricting the number of sites that can be sampled within a given sampling day. eDNA doesn’t face similar challenges and thus has the potential to greatly expand the ability of managers to monitor the ecosystems they manage.

Comparison with Reef Check visual surveys demonstrated both strong concordance between methods and important differences in the type and breadth of information they provide. After excluding *Sebastes*, eDNA detected an average of 83.5% of the taxa observed by Reef Check for the same sites during the same month and year. Some of this discrepancy may result from eDNA and Reef Check surveys not being conducted at the same exact time. The two approaches also differed substantially in the breadth of species detected with eDNA detecting far more species than Reef Check. This difference reflects, in part, the distinct capabilities of the two approaches. Reef Check surveys a predefined set of targeted species and provides observations on abundance, body size, and biomass. In contrast, eDNA surveys a much broader set of taxa but can only provide presence/absence data. The two approaches therefore provide complementary information that can help inform the success of MPAs. Our study contributes to a growing literature^20,44,45^ highlighting the value of pairing visual surveys with eDNA to leverage complementary strengths.

In contrast to visual based methods, field collection of environmental DNA is logistically simple and requires minimal effort, training, and equipment. In our sampling design, the primary requirement for field work was the ability to access the site and collect a liter of water while standing at knee depth. This low barrier to entry enabled participation across diverse age groups, demographics, and previous levels of exposure to science. Similar models of community science have been implemented into other large-scale eDNA projects, such as CALeDNA^46^. Beyond eDNA allowing community scientists to expand the scale at which samples can be taken, eDNA is also more cost-effective. Previous work in Southern California MPAs estimated that eDNA surveys were nearly 60% less costly than conventional scuba-based monitoring^20^. Together, using both eDNA and community scientists substantially increases sampling ability while reducing cost and effort compared to traditional methods.

Equally important is the value that community science extends beyond its benefit to research. Community participation in scientific research has been shown to enhance scientific literacy^47^, lead to more informed conservation advocacy^48^, create a sense of ownership/connection with the natural environment^49^, build networks of like-minded individuals^50^, incorporate local knowledge into scientific expertise^51^ and encourage engagement in local decision-making processes^47^. Specifically in the context of MPAs, where community buy-in and compliance with rules are important for long-term sustainability, community involvement in biodiversity may help to derive benefits beyond just monitoring such as greater engagement of communities in local conservation activities and a deeper connection to biodiversity.

### 4.3 Limited inference of MPA impact from site-to-site comparisons underscores the importance of long-term, within-site eDNA monitoring for MPA effectiveness

To evaluate the impact of marine protected area designation on nearshore fish assemblages, we compared two protected sites against two nearby non-protected comparison sites. At the species level, our data shows little differences between the MPAs and the non-MPAs. While just at or below the threshold for statistical significance (p = 0.052 and 0.048), alpha diversity analyses showed that the non-protected comparison sites recorded higher observed species richness than the protected areas. Similarly, comparative analyses of eDNA index values (i.e. relative abundances) found only one taxon with significantly greater relative abundances within MPAs compared to their paired comparison sites. In contrast, a small number of species at the SMR comparison and SMCA comparison sites showed higher relative abundances.

While this result may seem paradoxical, previous studies have observed similar results^20^. One possible explanation is the differences in size and biomass of fish within protected sites compared to non-protected sites. Reduced fishing pressures from MPA protections allow fish to increase in size^52^. Greater biomass would allow for common fish species to contribute disproportionately more DNA^53^ and comprise a larger fraction of total reads within the MPA’s samples. Conversely, in the non-protected areas with smaller and less abundant common species, less common taxa may make up a larger proportion of reads in the eDNA samples. This would lead to a higher observed richness or relative abundance in non-protected sites compared to protected sites. Additionally, species richness is just one indicator for MPA impact. Many of the ecological responses that are associated with MPAs, particularly in the Southern California MPA network, are increases in size and biomass, characteristics that eDNA metabarcoding does not directly quantify.

In contrast to alpha diversity patterns, beta diversity analyses revealed significant differences in the community composition of each site. Site-level community composition differed significantly among protection status and sites, suggesting that MPA status may be impacting community composition dynamics that are only detectable in multivariate space. However, the proportion of variance explained by seasonal and interannual variation explained the same or greater variation than site alone, indicating that protection status may not be driving changes. This is further complicated by differences in dispersion in the SMR site compared to the comparison site. When looking at the CAP ordination plots, we see distinct clustering of sites. Site characteristics such as microhabitats, geographical features, and localized conditions may be exerting a stronger impact on community structuring than MPA status alone.

Several additional factors may have contributed to the limited observed impact of MPAs. Studies show that both larger and older MPAs are positively correlated with MPA impacts^1,54^. First, the Point Dume SMR and SMCA are two relatively small protected areas at 7.5 square miles and 15.92 square miles respectively. Within these two sites, the habitat size themselves are also relatively small, with rocky reefs covering just 0.5 and 0.33 square miles at the SMR and SMCA^55^. Second, these two MPAs are relatively young. At the time of sampling, the MPAs had existed for 7-9 years after their implementation in 2012. For many fish species, research suggests that the timescales needed to observe a detectable difference due to MPAs is >10 years^56–58^. Therefore, one might not yet expect to see changes in fish communities in these MPAs.

These findings underscore a key limitation of traditional MPA monitoring. In dynamic coastal systems with localized habitat conditions impacted by broader global climatic drivers, paired MPA-“control” site comparisons may have minimal utility in evaluating whether MPAs are effective. This is a major challenge to inside-outside MPA comparisons broadly, especially in the California MPA network, as many sites like Point Dume SMR have no obvious analogous habitat and were intentionally chosen given their high biodiversity and habitat integrity. Furthermore, using sites that were chosen during baseline monitoring over a decade ago may no longer be adequate ecological equivalences given changes in erosion, ocean dynamics, and other abiotic forcings. For these reasons, long-term, within-site monitoring of MPAs may provide a more powerful framework for evaluating MPA effectiveness than site-to-site comparisons with “controls”. By establishing a baseline for each site and then doing yearly tracking within the MPA for species richness, relative abundances, and community composition, MPA effectiveness can be compared against itself. Comparison sites would remain important when not interpreted as controls for MPA success but as long-term sentinel sites that capture how regional biodiversity is responding under shared climate conditions^59^. Long-term monitoring at non-protected sites helps distinguish MPA-specific patterns from border environmental variability and allows managers to better contextualize whether observed changes inside MPAs reflect protection status or regional climate impacts. Given its scalability and cost-effectiveness, eDNA is particularly well suited for generating the high-frequency longitudinal time series needed for such monitoring.

### 4.4 Designing a low-cost, optimized eDNA sampling design for California’s 124 MPAs to capture nearshore fish diversity with substantially reduced field effort

A key objective of this study is to inform empirically grounded recommendations for an optimized eDNA sampling design for a long-term, within-site statewide MPA monitoring program. We aimed to identify the thresholds that balances coverage of biodiversity at the site with operational feasibility for a statewide program. To accomplish this, we used multiple analyses to identify the minimum number of sampling dates, samples, and sequences needed to capture 95% of the species diversity at a site.

Rarefaction analysis revealed that nearshore fish assemblages in Southern California require substantial sequencing depth. At the individual sample level, sequencing depth between ∼700,000 - 990,000 reads were needed to recover 95% of observed species richness in a sample.. Across the four Malibu sites, rarefaction analysis indicated that ∼11-14 samples were sufficient to capture a site’s species richness. To contextualize these thresholds into a monitoring framework, we conducted re-sampling simulations using the full two-year dataset to evaluate how sampling frequency influences total species recovery. With three biological replicates per site, quarterly sampling would have recovered approximately 95% of the total species observed over the study period, compared to monthly sampling. Collectively, these analyses indicate that moderate sampling frequency paired with a high biological sample replication of each site and a deeper sequencing depth can capture the majority of nearshore fish diversity that a monthly effort would. Notably, these findings diverge from other marine eDNA studies which report a much lower sequencing depth used. Previous Southern California eDNA surveys of MPAs found that species richness plateaued at a sequence sample size of 1,000 to 100,000 reads ^20^. While this study indicates that the read counts may need to be considerably higher than previously reported, advances in sequencing platforms are exponentially increasing the possible sequencing depths while reducing the costs per sample making this easily achievable.

Using this information, we present a rough base estimate for implementing a statewide eDNA MPA monitoring program. Based on inflation-adjusted supply costs, a workflow designed to achieve the sample and sequencing threshold, and the salary of two full-time state Environmental Scientist for analysis, we estimate quarterly sampling and analysis of all 124 MPAs, and one comparison site per MPA, would cost ∼$310,000 (Supplemental Table 4). While this estimate includes field consumables, lab work, and data analysis it does not include things such as the incidental costs, managing volunteer participation, shipping samples, or supplemental primers sets beyond MiFish. For comparison, currently advertised commercial eDNA metabarcoding services across companies generally cost $114-$145 per sample for laboratory processing, sequencing, bioinformatic analysis, and data delivery or $155 per sample including collection materials. At 2,976 samples in this study, that would correspond to ∼$461,000 per year for outsourced processing, although this price may not support the same sequencing depth needed. Therefore, this figure should be interpreted as a baseline estimate for eDNA monitoring and as a comparative figure for the potential cost of statewide monitoring.

An important assumption of this cost estimate is that eDNA sampling could be integrated into existing community science or monitoring networks. Our study benefited from an established partnership with Heal the Bay and an active local volunteer network, which reduced the logistical burden associated with recruiting and training participants. Comparable community science capacity cannot be assumed across all 124 California MPAs. Existing programs such as MPA Watch, Reef Check California, MARINe, PISCO, and other local or regional monitoring efforts provide the potential infrastructure to incorporate eDNA sampling, but their geographic coverage and volunteer capacity vary across the state. MPAs without an established community science network in the area would require additional investment in program coordination. However, a hybrid model where eDNA is incorporated into these pre-established networks of volunteers, while new partnerships or sampling capacity are supported in other areas, could provide an easy yet powerful long-term monitoring program for California MPAs.

While our study provides a strong case study for the value of eDNA in monitoring MPAs, extrapolating the values found here relies on a handful of assumptions across all MPA sites. The first is that this assumes all sites would have the same sampling and sequencing needs. Sites with higher biodiversity require a greater number of samples^60^ or a higher sequencing depth, while less diverse sites may require the opposite. The thresholds presented here are also optimized for capturing the greatest number of species. While this study found there are no seasonal patterns in diversity, other eDNA surveys of Southern California environments, particularly seagrass beds, showed that fish assemblages are seasonally impacted^61^, which would warrant different sampling designs. Similarly, although this sampling design recovered approximately 95% of the diversity observed at our sites, monitoring strategies in other regions may need to be tailored to specific management targets. For example, if managers are interested in a highly migratory species that is only present seasonally, sampling should be timed to coincide with the expected occurrence.

Beyond monitoring the impact of MPAs, a network of statewide eDNA sampling regimes would also provide considerable value to other management concerns. The eDNA samples could be analyzed with multiple primers to measure for plankton, invertebrates, marine mammals, etc. depending on concern. It could also be used to monitor for harmful algal bloom species^62^ or for range and distribution of invasive species^63^. Archived samples would also provide the ability for future reanalysis for things like tracking species shifts over time^64^. The multiplicative value of eDNA increases the value that a program like this would contribute and supports increased coastal monitoring with minimal increases to budget.

## Data Availability

All analyses were conducted in R and made available here: https://github.com/tannerwaters/MPA_study

The raw sequences generated for this project are available on the NCBI Sequence Read Archive under BioProject #PRJNA1531947.

## Acknowledgements

We thank the over seventy community scientists for helping collect eDNA samples in the field and spending their precious free time learning about California marine biodiversity and marine protected areas. We thank Elizabeth Reid-Wainscoat for supporting field collection efforts.

## Funding

TW was supported by the Eugene-Cota Robles fellowship from UCLA, the National Science Foundation Graduate Student Fellowship, the Qiagen Young Scientists Research Award, and the Switzer Foundation Fellowship. TW, OS, and LG were also supported by the Center for Developing Leadership in Science, which is supported by National Science Foundation grants (RISE-2024426 and RISE-228198), the Waverley Street Foundation, and by gifts from Oceankind and Dalio Philanthropies. Sequencing was carried out at the DNA Technologies and Expression Analysis Cores at the UC Davis Genome Center, which is supported by NIH Shared Instrumentation Grant 1S10OD010786-01. Experimental costs were supported by a gift from the Albert and Elaine Borchard Foundation, the Freedom Together Foundation, and grants from the David and Lucile Packard Foundation (2020-70931; 2022-74729) to RE. ZG was supported by NOAA’s Pacific Marine Environmental Laboratory and the NOAA’s Omics Program, contribution number 5879. ZG was also supported by a National Science Foundation Graduate Student Fellowship. Additional funding was provided by the UCLA La Kretz Center for California Conservation Science, UCLA Ecology and Evolutionary Biology Department, and National Geographic Explorers Grant to ZG and PB. The funders had no role in study design, data collection and analysis, decision to publish, or preparation of the manuscript.

## Author Contributions

Conceptualization: Gold Z, Parker E, Barber P; Methodology: Waters T, Gold Z, Parker E; Investigation: Waters T, Gold Z, Parker E, Simon O, Jenny L; Formal Analysis: Waters T, Gold Z, Parker S, Simon O, Jenny L; Visualization: Waters T, Simon O, Jenny L; Writing - original draft: Waters T; Funding Acquisition: Waters T, Eagle R; Supervision: Ambrose R, Barber P, Eagle R. Writing - editing draft: all authors.

## Competing Interests

The authors declare no competing interests.

## Supplemental Figures and Tables

**S1 Fig.**
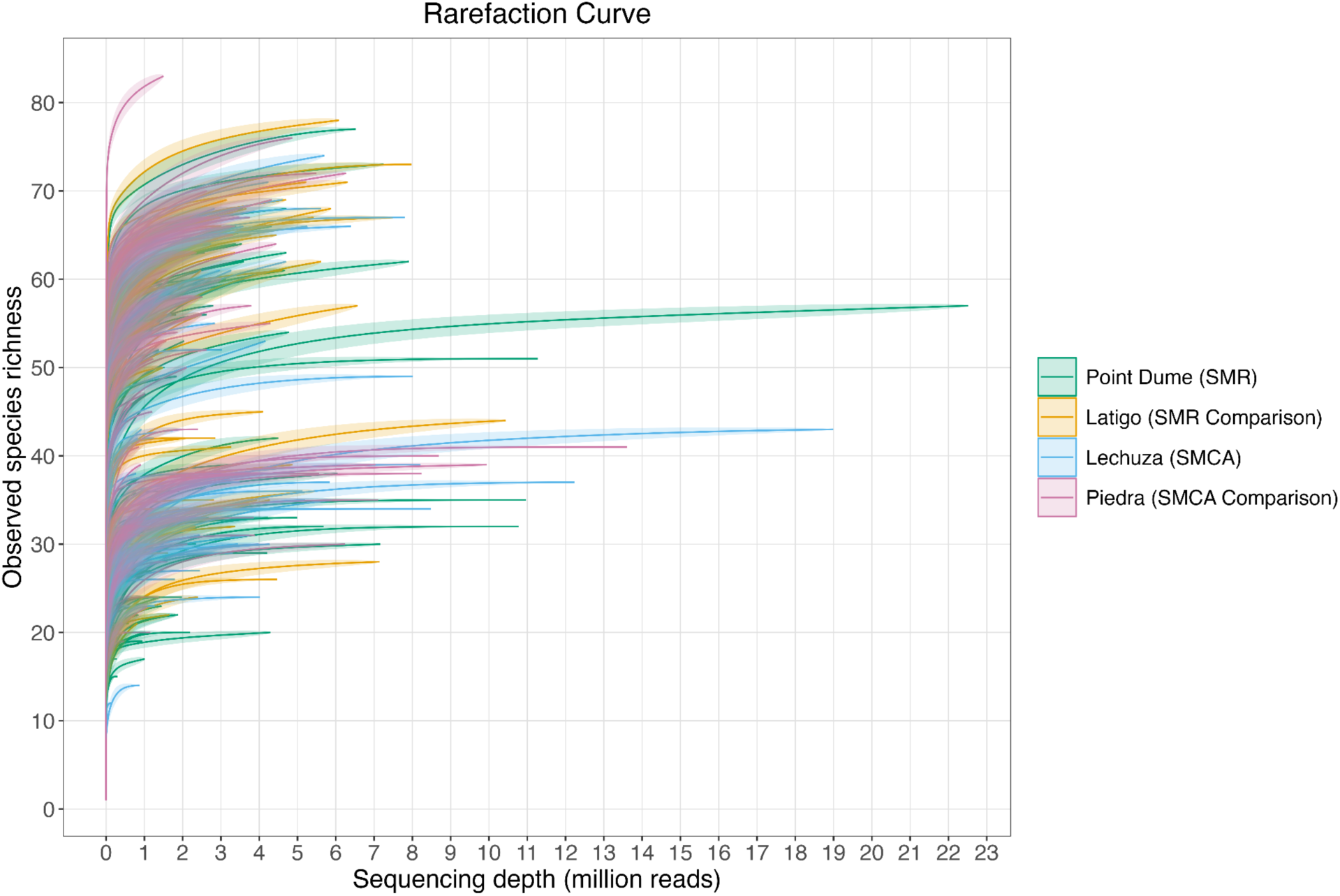
Sample rarefaction curves by site.

**S2 Fig:**
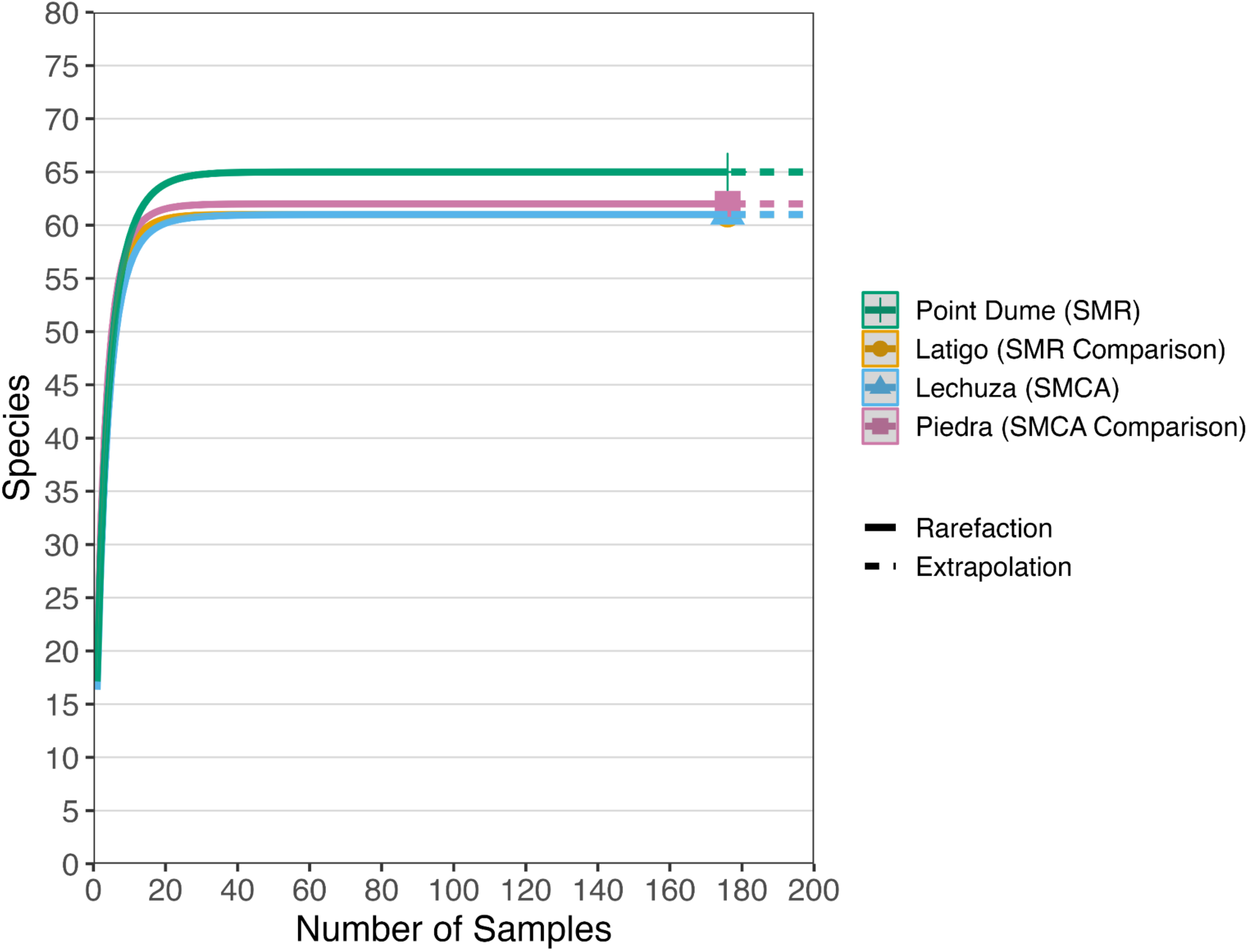
iNext based site-specific rarefaction curve showing species richness vs samples taken.

**S3 Fig.**
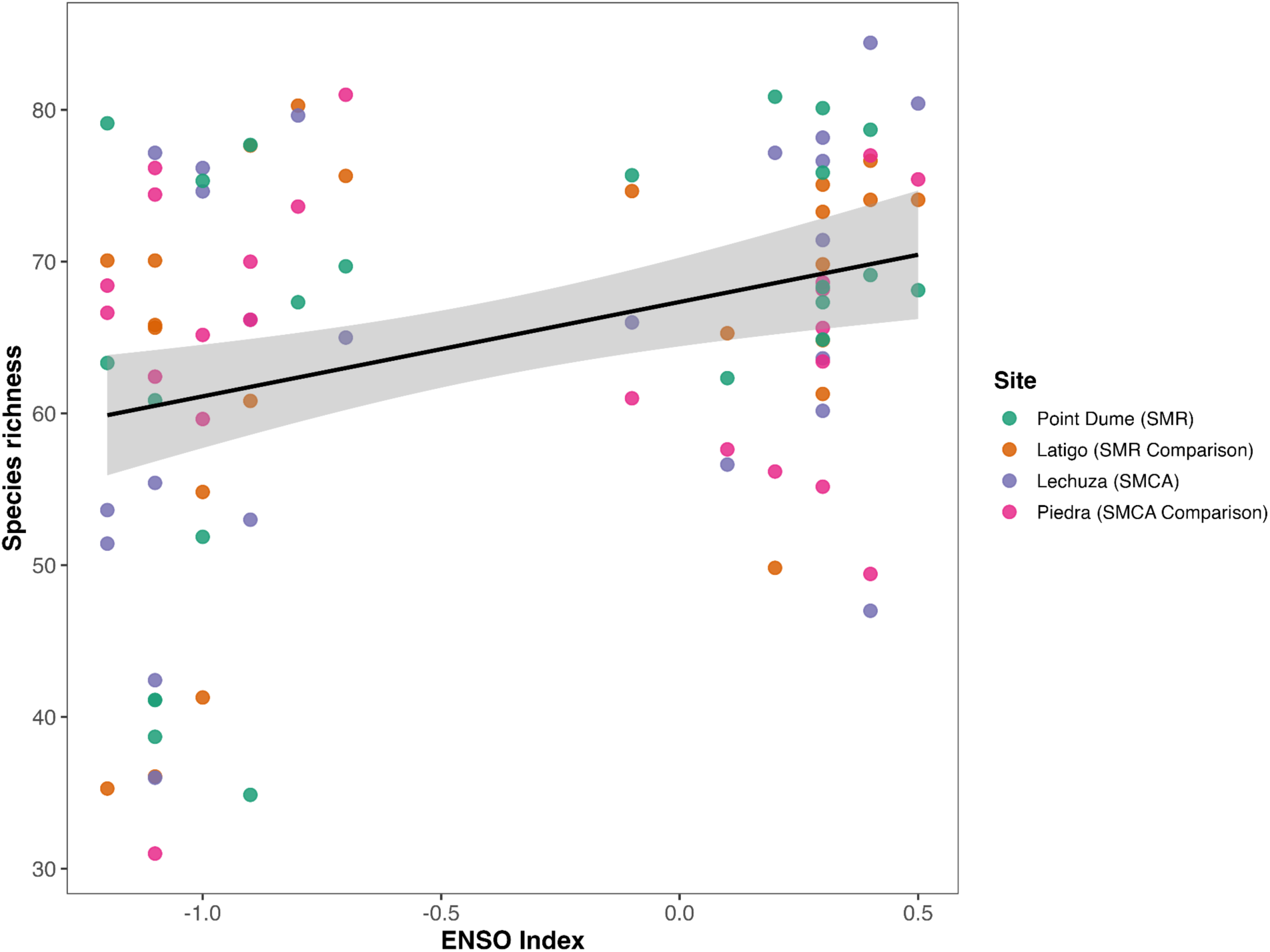
Monthly species richness was positively associated with the ENSO index after accounting for site and season. Each 1-unit increase in the ENSO index is associated with 6.25 additional species (p = 0.0023, R^2^= 0.211).

**S4 Fig.**
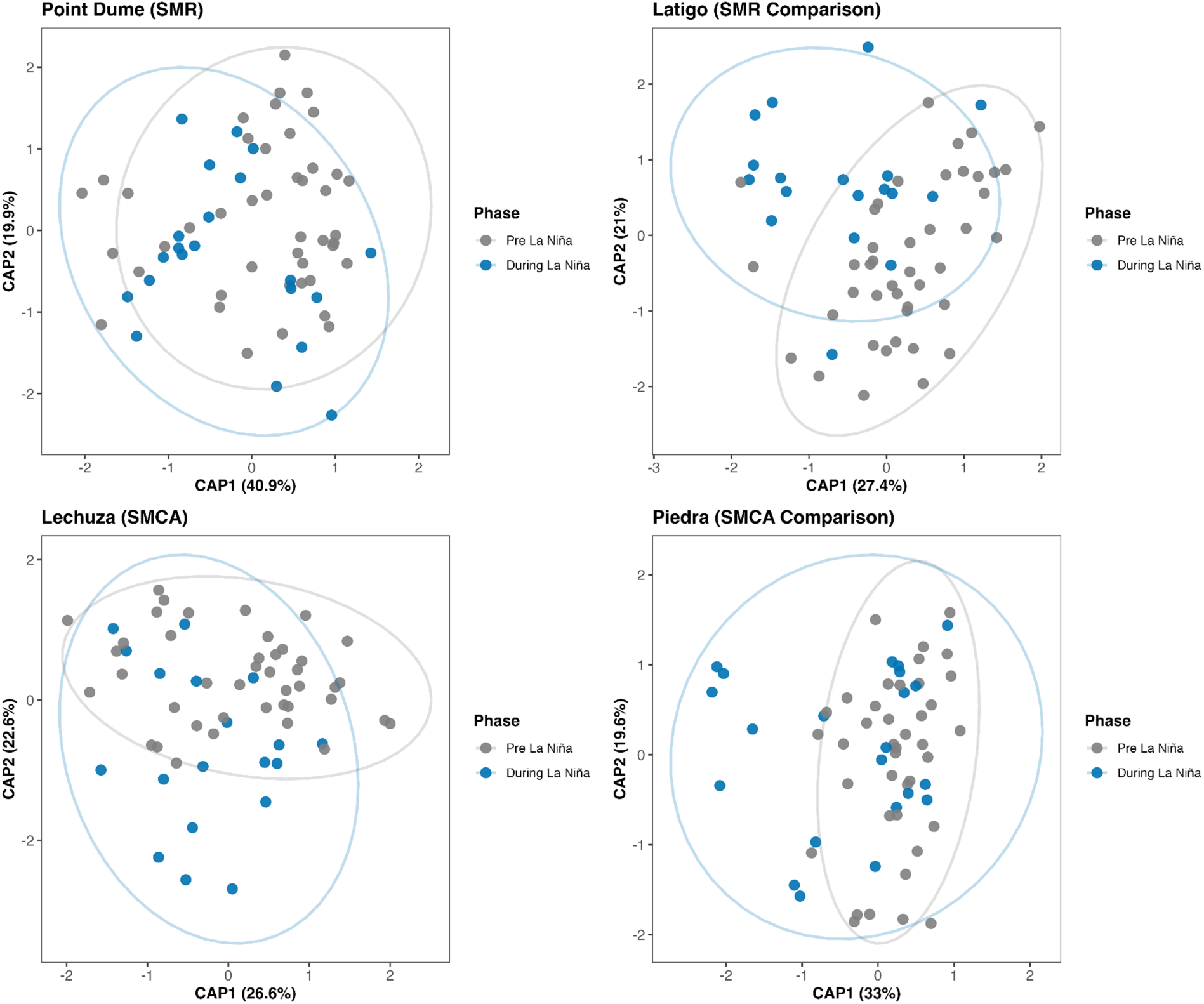
CAP ordination of community composition comparing pre-La Niña and during-La Niña periods across the four sampling sites. Points represent monthly sampling and ellipses indicate 95% confidence intervals. Most site-level differences between periods were modest with seasonal variation explaining more compositional variance than the La Niña phase.

## Notes

### Competing Interest Statement

The authors have declared no competing interest.

